# Persistent homology: a tool to universally measure plant morphologies across organs and scales

**DOI:** 10.1101/104141

**Authors:** Mao Li, Margaret H. Frank, Viktoriya Coneva, Washington Mio, Christopher N. Topp, Daniel H. Chitwood

**Affiliations:** Donald Danforth Plant Science Center, St. Louis, MO USA; Department of Mathematics, Florida State University, Tallahassee, FL, USA; Independent Researcher, St. Louis, MO USA

## Abstract

Genetic contributions to plant morphology are not partitioned between shoots and roots. Yet, shoot and root architectures are rarely measured in the same plants. Even if shoot and root architectures are both studied, the application of mathematical methods flexible enough to accommodate the disparate topologies and shapes within a plant, and across scales, are lacking. Here, we advocate the use of persistent homology, a mathematical method robust to noise, invariant with respect to orientation, capable of application across diverse scales, and importantly, compatible with diverse functions to quantify disparate plant morphologies, architectures, and textures. To demonstrate the usefulness of this method, we apply persistent homology approaches to the shape of leaves, serrations, and root architecture as measured in the same plants of a domesticated tomato Solanum pennellii near-isogenic introgression line population under field conditions. We find that genetic contributions to morphology affect the plant in a concerted fashion, affecting both the shoot and root, revealing a pleiotropic basis to natural variation in tomato.

## Introduction

Plant morphology cannot be partitioned; every part of the plant contributes to its morphology as much as any other (Chitwood and Topp, 2015; Topp et al., 2016). Yet, when measuring morphological traits, plants are routinely discretized, especially into above- and belowground parts. Any model predicting phenotype from genotype would be incomplete without considering the shoot and root as a whole. If morphology truly imparts function, then because of the extensive molecular, metabolic, hormonal, and physiological intercommunication between the root and shoot (Molnar et al., 2010; Thieme et al., 2015; Albacete et al., 2015; Warchefsky et al., 2016), even slight morphological changes in one would impact the other. The development of the shoot and root are also governed by similar molecular pathways (Wysocka-Diller et al., 2000; Sarkar et al., 2007; Stahl et al.,2009; Carlsbecker et al., 2010; Slewinski et al., 2012), which would suggest genetic changes affecting either could potentially impact both.

Even if root and shoot morphology are measured in the same plants, the different architectures and accessibilities of each has, until recently, necessitated disparate types of analyses. Roots and branches are intrinsically topological, and have been measured using a variety of length and width measures, branching parameters, skeletonization, shape descriptors, network properties, and other summary statistics (e.g. Drew, 1975; Fitter, 1987; Berntson, 1996; Lynch et al., 1997; Iyer-Pascuzzi et al., 2011; Bucksch et al., 2014). Contrastingly, many shoot organs can be represented as shapes, including leaves and other lateral organs (such as petals), fruits, and seeds. Unlike roots and branches, these structures can be quantified using geometric morphometric approaches, such as homologous landmarks (Viscosi and Cardini, 2011; Hasson et al., 2011; Klingenberg et al., 2012; Wang et al., 2014; Chitwood et al., 2014a; 2016a; 2016b), pseudo-landmarks (Langlade et al., 2005; Bensmihen et al., 2008; Weight et al., 2008; Feng et al., 2009; Cui et al., 2010; Costa et al., 2012), Elliptical Fourier Descriptors (Kuhl and Giardina, 1982; Iwata et al., 1998; Iwata and Ukai, 2002; Chitwood et al., 2012a; 2012b; 2012c; 2013; 2014b; Chitwood, 2014; Iwata et al., 2015), as well as using a variety of shape descriptors (Gonzalo et al., 2009). Shapes without orientation, such as pavement cells, require other approaches, such as isolating individual lobes from convex hulls (Wu et al., 2016), and overall dissection of closed contours can be measured using methods such as bending energy (Backhaus et al., 2010; Kuwabara et al., 2011).

The use of different analytic methods, applied to different organs and scales, should be disconcerting. If the organ systems of a plant represent the inextricably linked morphological manifestation of information encoded by the genome, analyzing the shoots and roots separately falsely dichotomizes plants. To understand the phenotypic representation of genomic information requires measuring morphology globally, across roots and shoots, using a comparable mathematical and statistical framework that can accommodate the disparate architectures, across scales, found within a single plant.

Here, we advocate the use of persistent homology as a method to quantify the topologies of root and shoot architectures and the shapes of leaves and other lateral organs. Persistent homology approaches are robust against noise, can be implemented in an orientation-invariant manner, accommodate the diverse scales found in plant structures, and most importantly, are flexible, employing varied functions to describe disparate topologies, shapes, and textures. As an example of the utility of persistent homology, we describe mapping the genetic basis of plant morphology in the roots and shoots of individual tomato plants from the Solanum pennellii introgression line (IL) population (Eshed and Zamir, 1995). We describe different functions applied through a persistent homology approach to quantify leaf shape, serrations, and root architecture. We find that the ILs with the most substantial morphological changes compared to the parent background (cv. M82) are affected in all examined structures. By measuring plant morphology in both the shoot and root using a common mathematical approach, insights into concerted genetic effects globally impacting plant architecture were uncovered that would have been missed using conventional techniques. We end by describing the importance of adopting mathematical frameworks, such as persistent homology, to accommodate comprehensive phenotypic descriptions of plant morphology and its potential to integrate seemingly discordant types of data in the plant sciences through an inherently topological, network-based perspective.

## Results

### A persistent homology primer

Persistent homology is a topological data analysis method that can be used to quantify complex shapes and construct informative summaries of data (Verri et al., 1993; Carlsson, 2009; Edelsbrunner and Harer, 2010). Results from methods other than persistent homology often vary based on the scale or parameters chosen for analysis. Choosing an “optimal” scale for analysis may be difficult, or impossible if features of the data are represented across scales. Thus, instead of analyzing a single scale, persistent homology tracks the evolution of features across all possible scales. In a persistent homology framework, the features are homology groups, recording the connectivity information within a topological space. Homology groups are of different orders, including H_0_ (zeroth order homology, path-connected components), H_1_ (first order homology, one-dimensional holes), or H_2_ (cavities or voids) (Hatcher, 2002). Persistent homology not only records features within each scale, but it also examines how features persist across scales, both the scale that features are generated (“born”) and disappear (“die”). To do so, it requires a filtration (a nested sequence of expanding shapes). This arises from a continuous function assigning values on the domain, which can be the shape itself, or a larger bounded region, such as a rectangle enclosing the shape, for example.

Below, we apply a persistent homology framework to diverse morphologies in the same tomato plants of a near-isogenic Solanum pennellii introgression line population (Eshed and Zamir, 1995), measuring leaf shape, leaf serrations, and root architecture, to demonstrate the versatility of this approach.

### Persistent homology and leaf shape

We quantify tomato leaflet shape (Fig. 1A) by studying the leaflet contour (Fig. 1B), which is a 2D point cloud comprised of contour pixels. We center and normalize the leaflet contour to the centroid size to focus only on shape. Since the data may be noisy and leaflets possess an orientation, we choose a method that is robust to noise and blind to orientation. To robustly represent the leaf contour, we use a Gaussian kernel density estimator (Fig. 1C), which can estimate the density directly from the data (Hwang et al., 1994). It is a smooth function that achieves higher values where there are concentrations of data, which in this case are contour pixels, highly concentrated in areas such as the tip of the leaflet, serrations, and lobes. The density estimator decays as it moves away from the data. Noise is usually very sparse, such that the function has smaller values around noise. To be invariant towards rotation, we study shapes falling in an annulus centered around the centroid of the leaflet contour (Fig. 1D). This can be achieved (Fig. 1E) by multiplying the density estimator (Fig. 1C) by the annulus kernel (a function that highlights and smoothens the annulus) (Fig. 1D). If we depict the function value as height, then the function is intuitively visualized as a set of ridges (Fig. 1F). As a plane moves from the highest function values to the lowest, we study the function values as superlevel sets (that is, the shape above the plane). The plane sometimes touches the peak of new ridge so that it generates a new connected component; alternatively, sometimes the ridges merge so that one connected component disappears. We use a persistence barcode (Fig. 1G) to record these changes: a bar, recording the scale at which the component is “born” and “dies”, represents each H0 connected component. For each leaflet contour, we compute 16 such persistence barcodes corresponding to 16 expanding annuli (Fig. 1B) to represent the shape of each leaflet.

**Figure 1:**
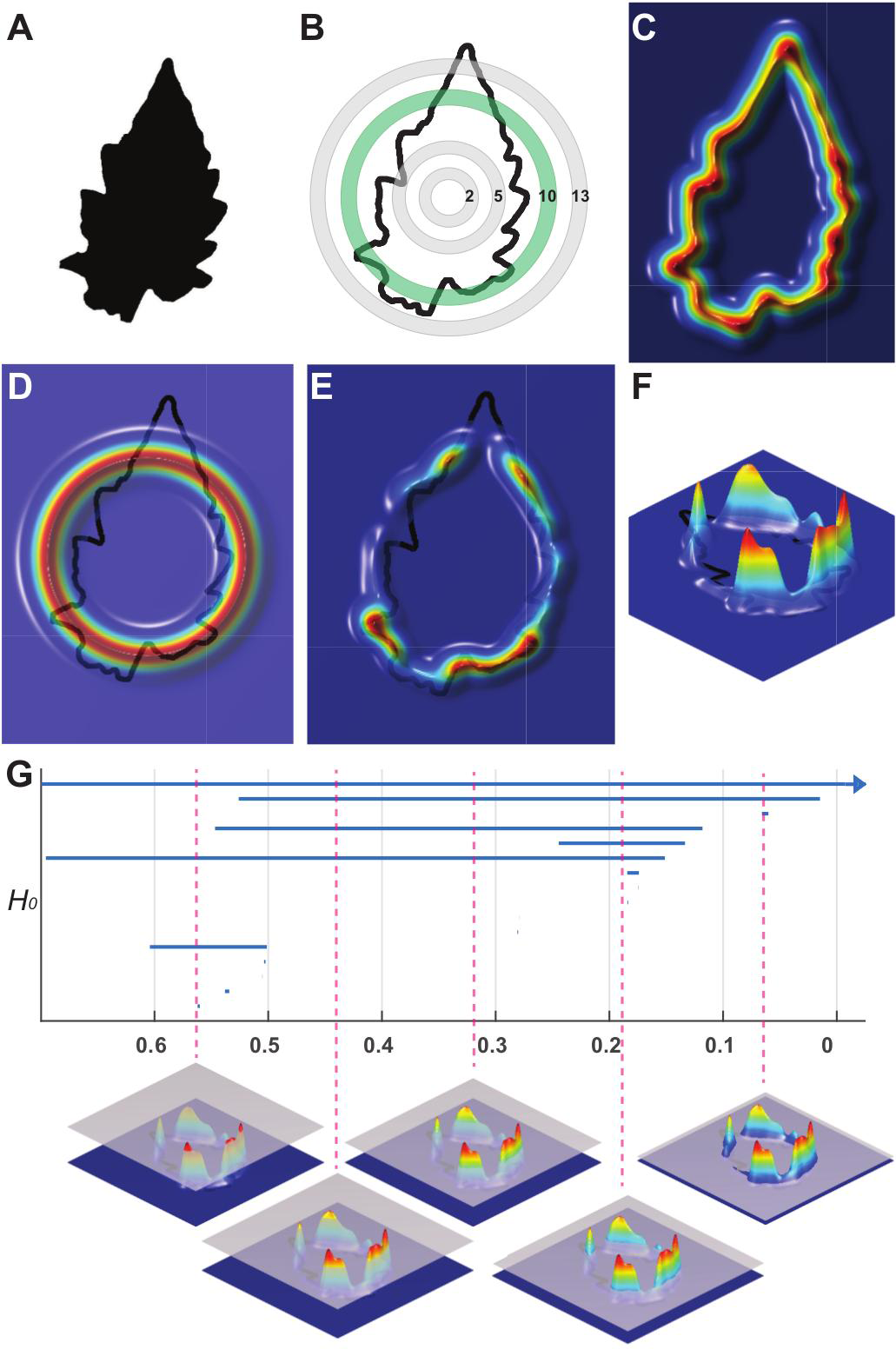
Persistent homology and leaf shape. **A)** A binary tomato leaflet image. **B)** Point cloud representing the contour of the leaflet and sampling of the 16 annuli used to subset data in analysis. The green ring (“10”) is used as an example subsequently. **C)** A colormap of a Gaussian density estimator applied to the contour point cloud. The density estimator is robust to noise. Red indicates a larger density of contour points (e.g., near serrations), blue a smaller density of contour points (e.g., straighter edges). **D)** An annulus kernel, a technique to localize and smoothen the isolation of density data. **E)** The multiplication of the density estimator (C) with the kernel annulus (D) emphasizes leaflet density features falling within the green ring indicated in (B). **F)** Side view of (E) showing the distribution of density features within the annulus. **G)** A persistence barcode. Each bar (vertical axis) represents the “birth” and “death” of a connected component. Connected components are “born” as different scales of the density features are traversed (horizontal axis) and “die” as they merge with other components. Eventually only a single component persists. The traversal of scales across density features is visually depicted (bottom), with magenta dotted lines indicating the relevant position in the persistence barcode (top).

### Persistent homology and leaf serrations

Although leaf shape contains some shape information about serrations, it is possible to craft persistent homology functions to focus on leaf serrations specifically. The serrations can be treated as the difference between the leaflet contour and a coarse approximation. To coarsely approximate the leaflet contour, we use Elliptical Fourier Descriptors (EFDs) (Kuhl and Giardina, 1982; Iwata et al., 1998; Iwata and Ukai, 2002). Such descriptors decompose the contour into a weighted sum of wave functions with different frequencies. Summing the higher frequency waves in the series, we describe finer details of the contour and achieve a closer approximation of leaflet shape. In contrast, if we use EFDs of the five lowest frequencies, we capture only coarse shape information (Fig. 2B). We then compute a distance function from the leaf contour to the EFD approximation. If the data point is inside of the approximated outline, we assign a negative sign to the distance value from the contour (blue), and if a data point falls outside, a positive sign distance value (red). We refer to this function as a signed distance function (Fig. 2C). Given a threshold (a number), the sublevel set is the points on the contour that have smaller values than the threshold. Changing the threshold continuously from small to large, we get the sublevel set filtration. Such sets can be roughly interpreted as the intersection between of the actual contour with a coarse shape of different sizes (the pink region in Fig. 2D intersecting the contour in dark magenta). We use an Euler characteristic (the number of connected components minus the number of loops) curve (Fig. 2D) of the signed distance function between the leaflet contour and the EFD approximation to quantify leaflet serrations.

**Figure 2:**
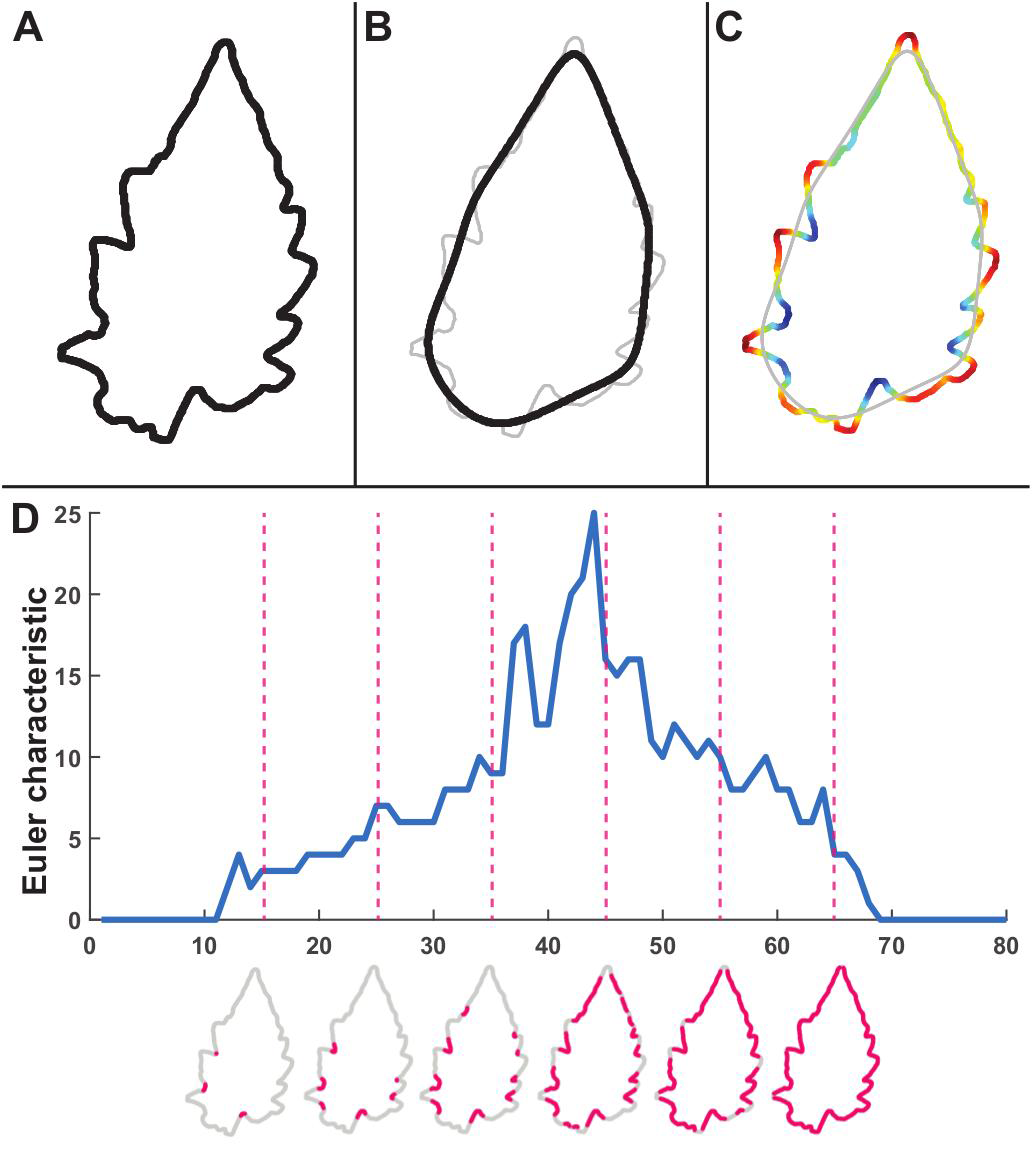
Persistent homology and leaf serrations. **A)** Point cloud representing the contour of a leaflet. **B)** A coarse approximation of the contour using an elliptical Fourier transform. **C)** A colormap of a signed distance function from the contour to the Fourier approximation. **D)** The Euler characteristic (the number of connected components minus the number of holes) curve. The Euler characteristic is plotted against traversal of the signed distance function. Shown are example leaves depicting the number of connected components (dark magenta) at the indicated positions in the curve (magenta dotted lines).

### Persistent homology and root architecture

To measure root architecture, we took a “shovel-omics” approach, in which roots of field-grown plants are dug up, washed, and photographed, capturing root architecture as a two-dimensional projection (Bucksch et al., 2014; Das et al., 2015) (Fig. 3A). The architectural complexity of a root is partly captured by the crossing of branches in the projection. Whereas traditional 2D root metrics are confounded by such crossings, we take advantage of them to quantify the complexity of the root system. When branches cross, they form loops, which is first-order homology (H1). We assign each pixel a value based on the distance to the root (Fig. 3B). Given a threshold (a distance value), we study the shape consisting of the pixels that have smaller values than this threshold. We record the number of loops (β1) formed by this shape. As we continuously vary this threshold, the number of loops also varies and becomes a β1 curve (Fig. 3C). We use this curve to quantify the complexity of branching pattern of 2D root projections.

**Figure 3:**
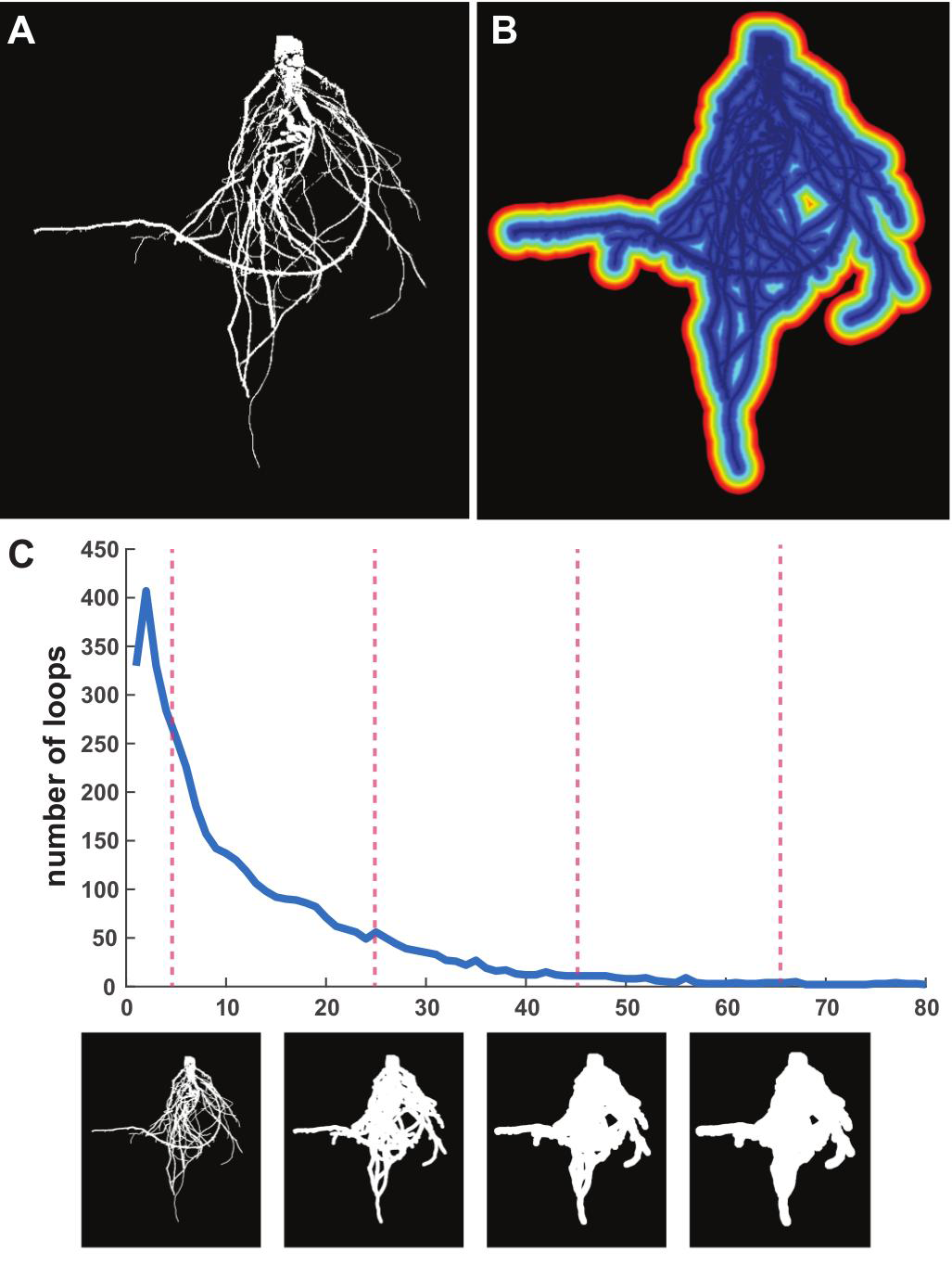
Persistent homology and root architecture. **A)** A binary image of root architecture as a 2D projection. **B)** A colormap of a distance function of pixels to the root. **C)** A β1 curve plotting the number of loops as a function of the distance function to quantify the complexity of root architecture. Shown are example root images (bottom) at indicated positions in the curve (magenta dotted lines).

### Persistent homology can detect global, genetic changes to shoot and root architectures

As described above, a persistent homology framework, combined with tailored functions, can describe morphologies as diverse as the shape of leaves (Fig. 1), serrations (Fig. 2), and root architecture (Fig. 3). By measuring these features in the same plants across a morphologically diverse population, we can determine the extent that genetic alterations globally or locally influence plant architecture. To do so, we leverage the near-isogenic Solanum pennellii tomato introgression lines (ILs) (Eshed and Zamir, 1995). These lines each harbor a single, relatively small introgressed region from the wild desert tomato S. pennellii in an otherwise domesticated tomato background of the cultivar M82. Detecting a significant phenotypic difference in an IL compared to the cv. M82 parent delimits the underlying genetic cause to the introgressed region (Chitwood et al., 2013). As we describe below, the ability to compare each IL against the cv. M82 parent provides a convenient means to reduce highly multivariate persistent homology data describing global features of complex morphologies into single discriminant values. These discriminant values can be correlated with each other to test hypotheses of how the morphology of different organs change in a concerted fashion or independently from each other in the presence of different introgressed regions. In the examples below we compare the introgression line IL4-3, which has large, previously documented changes in leaf shape and serrations (Chitwood et al., 2013; 2014b), to the cv. M82 parent.

We use 16 persistence barcodes, derived from 16 rings (Fig. 1), to quantify each leaf shape. For each pair of leaf shapes, we compute the bottleneck 2-distance (compute the bottleneck distance for each ring, then take the square root of the sum of these 16 bottleneck distances squared) between them. From all pairwise distances, we employ multidimensional scaling (MDS) to project the data into a reduced dimensional Euclidean space, to preserve the pairwise distances as much as possible. We then apply a canonical variant analysis (CVA) to the reduced dimensional projection to determine the discriminant features for two groups, which in this case is each individual IL compared to the cv. M82 parent. Because there are only two groups (IL vs. cv. M82), there is only one canonical variant (CV1). For example, CV1 values separating IL4-3 and cv. M82 is a composite of persistent homology features discriminating these two groups (Fig. 4A). A bootstrap test demonstrates that the detected difference between the mean CV1 scores of these two groups has high statistical significance (p<0.0001; Fig. 4). To interpret the morphological differences between lines, we can compute the contributions for each annulus using a linear discriminant analysis. For each ring, we compute the ratio of between-class scatter to within-class scatter to measure the contribution of each ring. For the comparison between IL4-3 and cv. M82, we see that the 16^th^ ring contributes the most discrimination between these genotypes (Fig. 4A). From the illustration (Fig. 4A), we can see that the difference is mainly derives from leaf length. The sixth and tenth rings contribute to the discrimination of the IL4-3 phenotype as well, and we can see that these differences mainly result from leaf width and lobing. The length and width of IL4-3 leaves has been previously shown to be some of the most discriminating features of this introgression line (Chitwood et al., 2013; 2014b).

**Figure 4:**
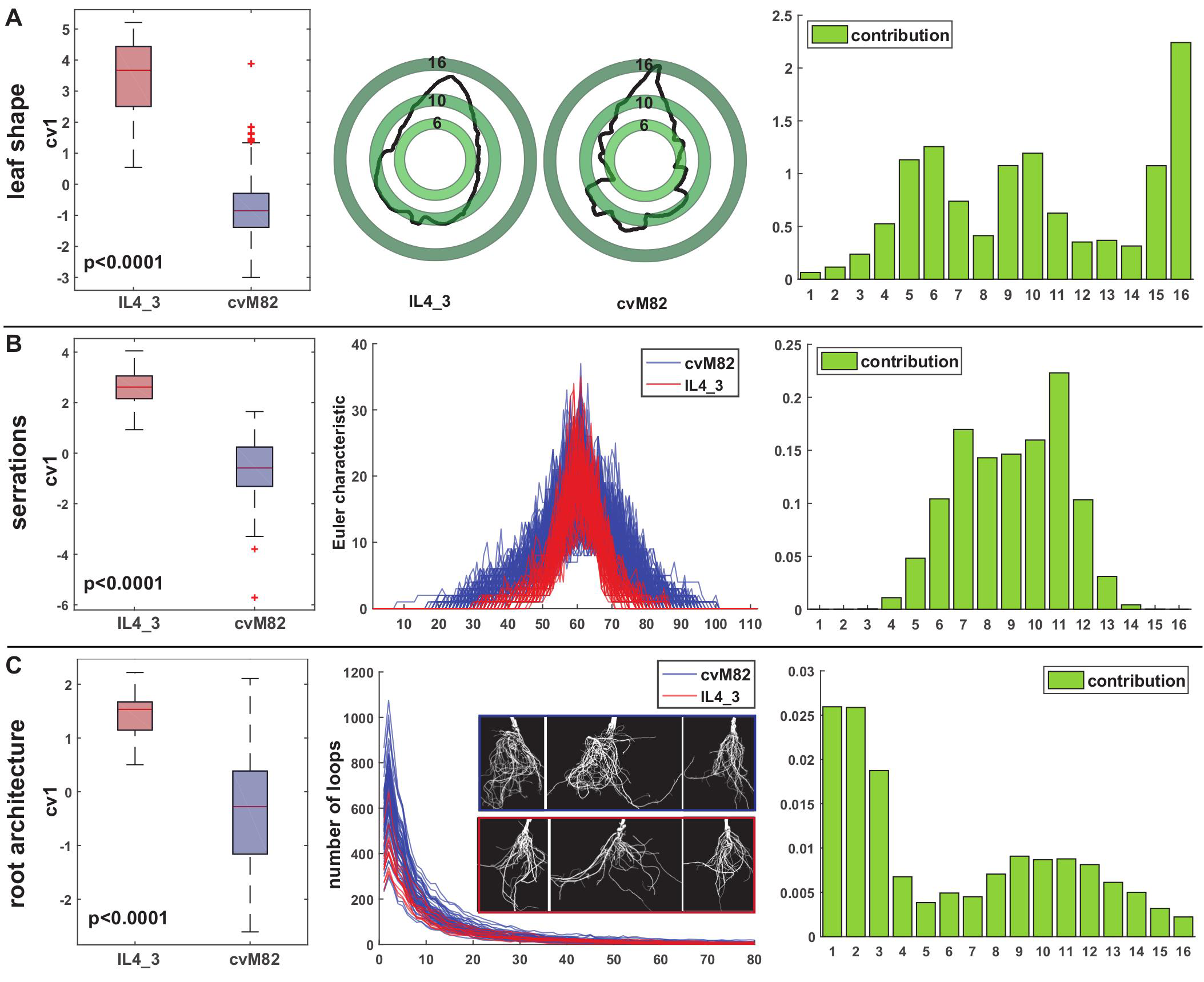
Comparing persistent homology results between IL4-3 and the parent cv. M82. **A)** Analysis of leaf shape. A boxplot of the canonical variant scores (CV1) between IL4-3 (red) and cv. M82 (blue), representing the discrimination of these two genotypes by 16 persistence barcodes (left). Example leaflets from each genotype and the superimposition of annuli used for analysis (middle). The contribution of rings to discriminating IL4-3 and cv. M82 (right). **B)** Analysis of serrations. The boxplot of CV1 scores (left), the Euler characteristic curves for all replications of IL4_3 (red) and cv. M82 (blue) (middle), and the contribution of every seven level sets of the Euler characteristic to discrimination of serration differences between the genotypes (right). **C)** Analysis of root architecture. The boxplot of CV1 scores (left), β1 curves depicting the number of loops across scalesfor all replications of IL4-3 (red) and cv. M82 (blue) and example root data (middle), and the contribution of every five level sets to root architecture discriminating IL4-3 and cv. M82 (right).

Another previously described feature of IL4-3 leaves is reduced serration compared to cv. M82, as quantified using circularity (a ratio of area to perimeter, 4π * 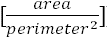)(Chitwood et al., 2013; 2014b). The Euler characteristic (EC) curves for all IL4-3 and cv. M82 replicates clearly indicate quantitative differences in leaflet serration as detected using persistent homology. Note that the Euler characteristic rises earlier and more quickly for cv. M82 (blue) compare to IL4-3 (red), indicating more numerous and deeper serrations (Fig. 4B). The EC curve for each leaflet is discretized into an 112-dimensional vector and then reduced in its dimensionality using a principal component analysis (PCA). Similar to leaf shape, we then apply a CVA and compute the CV1 score discriminating the genotypes by serration (Fig. 4B). Multiplying the PCA loadings by CVA loadings, we can measure the contribution of every seven vectors to the morphological differences between IL4-3 and cv. M82.

The first order betti number (β_1_) curves (the number of loops) for all replicates of IL4-3 and cv. M82 reveal differences in root architecture between these genotypes (Fig. 4C). We discretize each curve into an 80-dimensional vector and reduce the dimensionality by PCA to compute CV1 scores (Fig. 4C). The contribution for every five vectors to the root architecture differences is computed in the same way as inthe leaf serration analysis. Examining the β_1_ curves, it is evident that cv. M82 has greater root architecture complexity compared to IL4-3.

Following the analyses comparing IL4-3 to cv. M82 (Fig. 4), we compare other ILs with cv. M82 similarly (Fig. 5). Because persistent homology can detect exquisite morphological and architectural differences in a highly multivariate space, comparing each IL to cv. M82 using a discriminant analysis provides an indicator of global morphological differences for each IL, while allowing such differences to be fundamentally different in their nature. In other words, overall differences in leaf shape (Fig. 5A), leaf serrations (Fig. 5B), and root architecture (Fig. 5C), can be represented as a magnitude value (CV1), regardless of the myriad potential ways such morphological differences manifest.

**Figure 5:**
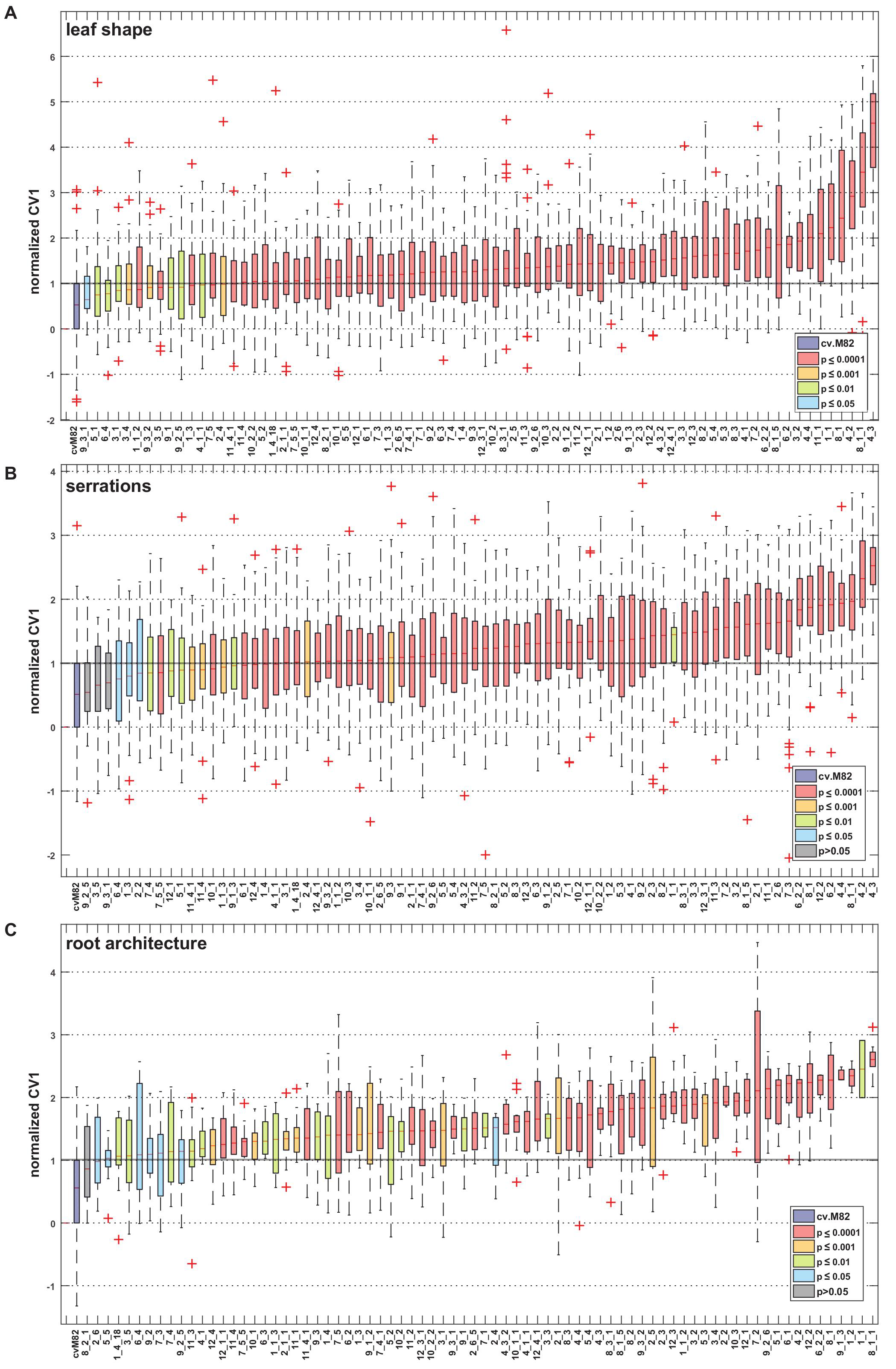
Overall morphological differences between near-isogenic introgression lines and the parent cv. M82. Normalized canonical variant scores (CV1) measuring global, discriminating differences between introgression lines (ILs) and cv. M82 plotted against IL identity, ranked by median CV1 value. Color indicates p values. Shown are CV1 scores for **A)** leaf shape, **B)** leaf serrations, and **C)** root architecture.

### Persistent homology detects concerted changes in shoot and root architecture

Applying a persistent homology framework to the S. pennellii ILs allows a single value encapsulating the global morphological features of different plant organs— leaf shape, serrations, and root architecture—to be calculated across genotypes. Such comprehensive traits differ from univariate traits that dramatically changed during domestication, such as fruit weight (Frary et al., 2000), for which a large amount of phenotypic variance is attributable to a single locus. Rather, persistent homology traits more closely resemble other multivariate traits, such as was used to identify QTL controlling 3D rice-root architecture (Topp et al., 2013), or the Fourier decomposition of leaf shape (Chitwood et al., 2013; 2014b), which is highly heritable but extremely polygenic. By reducing the myriad possible architectural changes in leaves and roots to a single value reflecting magnitude through discriminant analyses between ILs and cv. M82, we ask and begin to answer fundamental questions in plant biology: Do genetic alterations affecting one part of the plant affect others? Generally, are genetic changes in shoot architecture accompanied by changes in the root (and vice versa), or are these organ systems under separate genetic controls?

Comparing CV1 values normalized to cv. M82 across traits, it becomes apparent that ILs with the most profound differences from cv. M82 for one trait, tend to be profoundly altered in others (Fig. 6A). Eight ILs (IL3-2, IL4-2, IL4-3, IL6-2-2, IL7-2, IL8-1, IL8-1-1, and IL8-1-5) are among the strongly altered ILs for leaf shape (in the top 12), leaf serrations (in the top 15), and root architecture (in the top 27). Furthermore, the median values of the normalized CV1 values for each trait are significantly correlated with each other (Fig. 6B), although leaf shape and serrations are more highly correlated with each than with root architecture.

**Figure 6:**
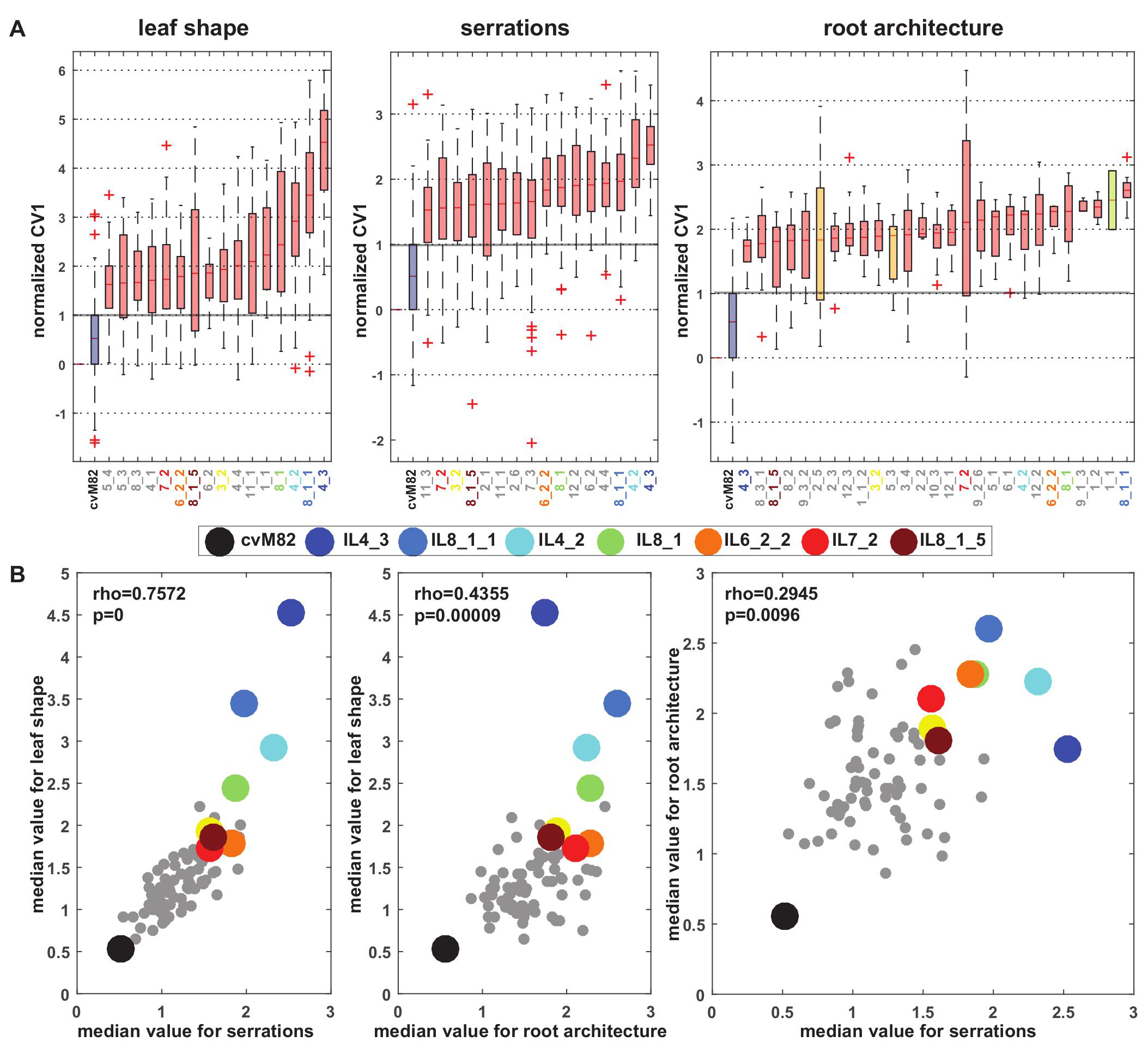
Concerted changes in leaf and root architecture underlie morphological diversity in tomato introgression lines. **A)** The most extreme ILs for each trait ranked by median CV1 value. Left, leaf shape; middle, serrations; right, root architecture. **B)** Scatter plots of median values for leaf shape vs. serrations (left), leaf shape vs. root architecture (middle), and root architecture vs. serrations (right). The most morphologically different ILs from cv. M82 are consistently so across traits. The most extreme ILs are similarly colored across all plots. Black indicates cv. M82.

Our results indicate that shoot and root architectures, when analyzed with a metric capable of quantifying multivariate features across scales, are under concerted genetic control. We emphasize that these changes only represent the magnitude of architectural changes, implying that changes in shoot morphology are simply accompanied by changes in root architecture (and vice versa), but that nothing is implied about the direction of the phenotypic changes within the multivariate space.

## Discussion

Persistent homology is a framework for analyzing topology at different resolutions, and as such bypasses major obstacles that currently confront the analysis of discretized shoot and root architectures. It can be applied in an orientation-independent fashion and by analyzing shape and topology through a continuum of resolutions, is robust to noise and accommodates features at multiple scales. The most versatile feature of a persistent homology framework is that any number of functions, specific to the task at hand, can be analyzed for topological properties at different resolutions. The examples of leaf shape, serrations, and root architecture in this work exemplify the utility of such an approach in the global analysis of plant morphology. Density functions applied through growing annuli originating from leaf centroids measure shape (Fig. 1); set levels emanating from a Fourier transform approximation of leaf shape measure serrations (Fig. 2); and dilation of root architectures can detect first order Betti number (β_1_) loops (Fig. 3). The metaphor of topology used across varying resolutions is an adaptable one, and when applied to different features throughout the same plants, can determine the genetic basis of global morphology (Figs. 4-5) and concerted changes in root and shoot architectures revealing previously undetected pleiotropy in tomato (Fig. 6).

It is worth contrasting persistent homology with conventional morphometric methods in plants. What persistent homology gains by being universally applicable it loses in being less intuitive, requiring an understanding of the functions used and how they are applied. For example, many conventional root measures are literal and easy to comprehend (such as root length and diameter, branching angles and densities, convex area, and other shape descriptors). But even in sum, they can represent only limited aspects of the morphospace, and thus our understanding of their genetic basis. Similarly, Procrustes-adjusted homologous landmarks and pseudo-landmarks have literal meaning and can be represented as eigenshapes, but the decision of which landmarks to use and the scale of the features they measure is also arbitrary. Even though somewhat abstract, Elliptical Fourier Descriptors (EFDs) can also be used to reconstruct eigen-representations of leaves but fail to measure a range of scales, sacrificing the local measurement of serrations for overall leaf shape. Features such as pavement cell lobes and leaf serrations and lobes can be isolated and measured for specific properties, but again, such approaches focus on local (and discretized) elements of plant morphology. Although straightforward to understand, all these approaches suffer from partitioning the plant into discrete units that do not exist in reality, capturing either magnitude or shape, or focus on only one scale of resolution, foregoing a truly comprehensive measure of morphology from local to global features.

By applying functions over a range of resolutions, persistent homology can measure local features in a global manner, and this is where its versatility and power lies. The multivariate representations of persistent homology results are therefore not an assembly of myriad, piecemeal measurements of parts of the plant defined by human intuition; rather, persistent homology results are represented as a continuum of function outputs applied over a range of resolutions. It is this underlying philosophy that allows persistent homology to be applied to any number of plant morphological features—topology, shape, texture (such as in pollen grains; Mander et al., 2013), and more—permitting an unprecedented, unified view of plant architecture. It will be particularly useful for analyzing dynamic changes in plant morphology over time, a reflection of the ability to apply persistent homology in n-dimensional spaces and of the requirement to analyze over ranges of resolutions. Importantly, persistent homology can be applied to networks, and by reducing datasets (such as gene expression, proteomic, metabolite, and other molecular data) to topologies, potentially represents a way to analyze disparate data types in a truly integrated fashion. Such a unified view of plant morphology (and plant science, in general) is desperately required in an age when the automated acquisition of plant morphological and molecular data is reaching unprecedented levels.

## Materials and Methods

### Plant material and growth conditions

*Solanum pennellii* introgression lines (ILs, LA4028-LA4103; Eshed and Zamir, 1995) and cv. M82 seeds were treated with 50% bleach for 1 min, rinsed with water, and germinated in phytatrays lined with moist paper towels. The seeds were left in the dark for 3 days, followed by 3 days in light, and transferred to greenhouse conditions in 50-plug trays. At this point, the plants were randomized according to a block design at a replication of 15. Plants were hardened by moving them outside for 10 days (5/10/2014). Hardened plants were transplanted to field conditions (5/21/2014, Bradford Research Station, Columbia, MO) with 10 feet between rows and 4-foot spacing between plants within rows. The final design had 15 blocks, each consisting of 4 rows with 20 plants per row. Each of the 76 ILs and 2 experimental cv. M82 plants were randomized within each block. After flowering (the week of 7/21/2014), four fully expanded adult leaves were harvested from each plant, and the adaxial surfaces of the left distal leaflets were scanned. The scans were processed using Image J macros to segment individual leaflets and to threshold and binarize each leaflet image. After fruit harvest, the roots of all plants were dug out manually, washed and imaged using a camera setup as detailed in Das et al. (2015).

### Persistent Homology

Persistent homology (Verri et al., 1993; Carlsson, 2009; Edelsbrunner and Harer, 2010) is a tool of topological data analysis that is well suited for the study of complex shapes such as plant morphological data extracted from digital images. Persistent homology applied to a function captures information about its shape by integrating topological signatures of its various sublevel or superlevel sets into a single persistence diagram. The data analyzed in this paper consist of 2D point clouds extracted from images that are converted into functions that encode local-to-global morphological properties such as leaf shape, leaf serrations, and root architecture. Our pipeline for converting 2D plant imaging data into persistence diagrams involves four main steps:

1. Extracting point clouds from images: From binary images, we extract 2D point clouds representing leaf contours or 2D projections of roots, where a pixel is converted into a point via its coordinates.
2. Converting point clouds to functions: From point cloud data, we construct various functions that carry local and global information about leaf shape, leaf serrations, or root architecture. A more detailed description of these functions is given below.
3. Forming filtrations: A filtration of a domain D is an expanding sequence of subsets that eventually cover D. In applications, D may be a leaf contour, a 2D projection of a root, or a full rectangle containing such data. A function on D yields a filtration as follows. Given a threshold value r (a number), the set of points in D whose function values do not exceed the threshold is called the sublevel set for r, which can be mathematically described as

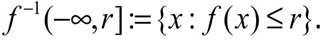

Similarly, the set consisting of points whose function values are not smaller than the threshold is called the superlevel set for r and given by

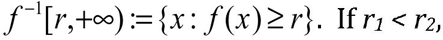

then the sublevel set for r1 is always included in the sublevel set for r2. As we reach the maximum value of the function, its sublevel set comprises the entire domain. Thus, we obtain a filtration called the sublevel set filtration. A similar construction applies to superlevel sets. Fig. 1G provides an example of a superlevel set filtration, where the domain D is a square containing a leaf. Fig. 2D depicts a filtration of the contour of a leaf in which sublevel sets correspond to intersections of the growing pink regions with the leaf contour. In Fig. 3C, the sublevel sets correspond to different dilations of a root.
4. Computing persistent homology: From a filtration, as described above, we construct barcodes that summarize the topology of the various stages of the filtration; more specifically, their 0-dimensional and 1-dimensional homology. Here, homology refers to a mathematical descriptor of the shape of the filtration, distinct from the concept of homology by descent from a common ancestor in biology. 0-homology captures the number of connected components (the number of islands) at each stage of the filtration. This number is known as the 0-th Betti number and is denoted *β*_0_. For example, in Fig. 2D, we see three connected components at the first stage, then six, etc. The full evolution as components are created or merge along the filtration can be encoded in a single barcode (Carlsson et al.,2005; Ghrist, 2008), as illustrated in Fig. 1G. A bar in a barcode starting at a value b (birth) and ending at d (death) indicates a connected component newly generated at the level b that merges with others at level d. Thus, more than just tracking the evolution of *β*_0_, a persistence diagram contains information about how components coalesce at different stages of the filtration. Similarly, 1-homology is about the number *β*_1_ of essential loops (holes), known as the first Betti number, leading to a barcode for the 1-homology of a filtration. There are higher dimensional analogues; however, they will not play a role in our are the data points on the contour and analyses because our data is 2-dimensional. We use the software package JavaPlex (Adams et al. 2014) to compute barcodes. Some useful reductions of persistence diagrams that are simpler to compute are the *β*_0_ curve, the
*β*_1_ curve, and the Euler characteristic curve, which are described next. As we vary the threshold r continuously, *β*_0_ also changes producing a *β*
_0_ curve that describes how the 0-th Betti number evolves with the threshold. Similarly, we obtain a *β*_1_ curve, as exemplified in Fig. 3C (to measure complexity of root architecture). For 2D domains, the Euler characteristic (EC) is given by

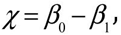
also viewed as a curve, as in Fig. 2D (to measure serrations). The Euler characteristic is the easiest to compute since it may be calculated directly from a triangulation of an object.

For statistical shape analysis based on persistence diagrams, a key ingredient is a metric to measure similarity between two persistence diagrams. The metric of choice is known as the bottleneck distance, with respect to which persistence diagrams are known to be robust features (Cohen-Steiner et al., 2007). We also carry out statistical analysis based on EC curves and *β*_1_ curves. These are Euclidean features that can be approached with standard techniques such as principal component analysis (PCA) and linear discriminant analysis (LDA).

A key step in the present approach to quantification of variation in morphology is the design of functions that effectively capture shape properties: local, regional, and global. Next, we describe the functions used for analyses of leaf shape, leaf serrations, and root architecture.

Leaf shape: Topological features often times become significantly more effective for shape representation and analysis under spatial localization. For leaf contour shape, we begin with a Gaussian density estimator for the entire contour of a leaf, defined as
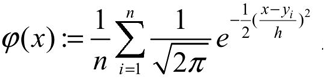
where *y*_1_, ◻, *y*_*n*_are the data points on the contour and h is a bandwidth parameter; see Fig. 1C. We then modulate this function by multiplying it by a “bump” function K that localizes it to a particular region. In our analyses, we use localization to concentric rings about the centroid of a leaf. The annular domains ensure that the resulting topological features are independent of orientation, as shown in Fig. 1D–Fig. 1F. More precisely, we use modulation by kernels of the form 
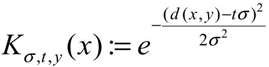
, where y is the center of the annulus, *tσ* determines its radius, and the parameter *σ* its width. We refer to persistent homology derived from such localized functions as local persistent homology.

Leaf serrations: Elliptical Fourier Descriptors (EFD) decompose a contour into a weighted sum of wave functions with different frequencies. Summing only over a finite number of harmonics gives a smooth approximation of the contour. This smoothing effect leads to loss in such details as serrations to a certain degree. However, we take advantage of this and quantify serrations by looking at residuals; that is, the difference between the original contour and the smooth approximation. Our experiments indicated that EFDs for the first five lowest frequencies yield smooth approximations suitable for serration analysis. Let C denote the original leaf contour and T the smooth approximation. We compute the distance from each point on C to T with the convention that if the point on C is inside the contour T, then we assign a negative sign to the distance. The distance is non-negative otherwise. We analyze serrations via the Euler characteristic function associated with the sublevel set filtration of this signed distance function. In this case, the EC curve is closely related to the*β*_0_ curve, but easier to calculate.

Root architecture: From an image of a 2D root projection, we construct a distance from function that computes the distance from a point to the nearest pixel on the root. In this way, all points on the root have value 0. The farther the point is from the root, the larger the value of the function. If we increase the threshold value starting from 0, the sublevel set filtration gives progressively larger dilations of the root. Since root branching typically creates numerous crossings and loops in 2D projections, we use the *β*_1_ curve associated with this distance function as a measure of complexity of root architecture.

All Matlab functions necessary to calculate persistence barcodes, bottleneck distances for leaf shape, euler characteristic curves for leaf serrations, and *β*_1_ curves for root architecture used in this manuscript can be found at the following GitHub repository:https://github.com/maoli0923/Persistent-Homology-Tomato-Leaf-Root

## Acknowledgements

The authors are indebted to Eric Floro and other members of the Topp Lab that contributed to fieldwork and root imaging.

## Supplemental data

All Matlab functions necessary to calculate persistence barcodes, bottleneck distances for leaf shape, euler characteristic curves for leaf serrations, and *β*_1_ curves for root architecture used in this manuscript can be found at the following GitHub repository: https://github.com/maoli0923/Persistent-Homology-Tomato-Leaf-Root

